# RNA pull-down-Confocal Nanoscanning (RP-CONA) detects quercetin as pri-miR-7/HuR interaction inhibitor that decreases α-Synuclein levels

**DOI:** 10.1101/2021.01.01.425030

**Authors:** Siran Zhu, Saul Rooney, Nhan T. Pham, Joanna Koszela, David Kelly, Manfred Auer, Gracjan Michlewski

## Abstract

RNA-protein interactions are central to all gene expression processes and contribute to variety of human diseases. Therapeutic approaches targeting RNA-protein interactions have shown promising effects on some diseases that are previously regarded as ‘incurable’. Here we developed a fluorescent on-bead screening platform: RNA pull-down-Confocal Nanoscanning (RP-CONA), to identify RNA-protein interaction modulators in eukaryotic cell extracts. Using RP-CONA, we identified small molecules that disrupt the interaction between HuR, an inhibitor of brain-enriched miR-7 biogenesis, and the conserved terminal loop of pri-miR-7-1. Importantly, miR-7’s primary target is an mRNA of α-Synuclein, which contributes to aetiology of Parkinson’s disease. Our method identified a natural product quercetin as a molecule able to upregulate cellular miR-7 levels and downregulate the expression of α-Synuclein. This opens up new therapeutic avenues towards treatment of Parkinson’s disease as well as provides novel methodology to search for RNA-protein interaction modulators.

## INTRODUCTION

RNA-protein interactions coordinate the whole of RNA metabolism, including transcription, RNA processing, modification, translation and turnover (1). Dysregulated RNA metabolism can result in serious diseases including cancer (2) and neuropathological conditions (3). In recent years, therapies targeting RNA-protein interactions have shown promising results and some of them are already in clinical trials (4–6). Among many types of RNAs, short non-coding microRNAs (miRNAs, miRs), that regulate gene expression by imperfect base-pairing to mRNA, hold great potential both as biomarkers (7) and therapeutics (8) in multiple human pathologies. The hairpin-like transcripts, known as primary miRNAs (pri-miRNAs), are processed by Microprocessor complex to become stem-loop precursor miRNAs (pre-miRNAs) (9–13). The intermediates are further excised by an RNase III enzyme Dicer, generating miRNA duplexes (14,15). Only the final product, the singlestrand mature miRNAs together with the Argonaute proteins are functional regulators of gene expression (16).

Parkinson’s disease (PD) is an incurable neurodegenerative disease that affects all ages but is most prevalent in the elderly population, with over 1% of those over the age of 60 suffering from this disease (17). One of the main causes behind PD is overproduction and aggregation of a protein called α-Synuclein (α-Syn), expressed from the SNCA gene in the brain cells of affected individuals (18–21). There is a large body of evidence that decreasing the levels of α-Syn should be beneficial for PD patients and several clinical trials are now focusing on α-Syn clearance with small molecules, antibodies or vaccines (22). Notably, miR-7 has been shown to target α-Syn production (23) and is significantly downregulated in the substantia nigra pars compacta (SNpc) of PD patients (24). MiR-7 also targets other genes implicated in PD, including RelA, Sir2 or Nlrp3 (25). For these reasons, approaches for miR-7 replacement therapies have been put forward (26).

The biogenesis of miRNAs is tightly controlled through RNA-protein interactions (27–31). Our group has identified that an RNA binding protein (RBP) HuR specifically binds to the conserved terminal loop (CTL) of primary miR-7 (pri-miR-7-1). By recruiting another RBP MSI2, HuR increases the rigidity of the pri-miR-7 stem loop, therefore, blocking Microprocessor cleavage and preventing production of mature miR-7 (32). HuR-mediated inhibition of miR-7 biogenesis was independently reported by Lebedeva et al. (33). We have also found that monounsaturated fatty acid - oleic acid (OA), previously found to bind to MSI2 (34), can dissociate the HuR/MSI2 complex from pri-miR-7 and facilitate the biogenesis of miR-7 (35). However, OA is not effective at micromolar concentrations, has poor bioavailability when tested in cells and is toxic in high concentrations (36). Thus, it is necessary to identify potent pri-miR-7/HuR inhibitors that will elevate miR-7 levels and (37)lead to downregulation of its targets, including α-Syn, which could provide alternative solutions to PD therapy.

Here we developed a fluorescent on-bead screening assay based on RNA pull-down Confocal Nanoscanning (RP-CONA), to identify small molecules that modulate the strength of RNA-protein interactions. Our method uses an ultra-sensitive RNA-protein pull-down assay in cell extracts from human cultured cells (38) detecting RNA-protein complex modulators by confocal microscopy (39–41). By employing RP-CONA, we identified quercetin, a natural flavonoid and known inhibitor of HuR/TNF-α mRNA interaction (42), as the most potent pri-miR-7/HuR interaction inhibitor. Quercetin induces miR-7 level and inhibits α-Syn expression in an HuR-dependent fashion, but the exact mechanism behind its activity remains to be fully elucidated. In summary, in this study we introduce RP-CONA as a new assay technique for the identification of small molecule modulators of RNA-protein interactions. We identified novel inhibitors of HuR/RNA complexes and have discovered a validated hit compound towards attenuation of α-Syn levels in PD.

## MATERIAL AND METHODS

### Cell culture

HeLa and HEK293T cells were maintained in DMEM (Dulbecco’s Modified Eagle Medium, Gibco) containing 10% foetal bovine serum (Gibco). Transfection was performed using Lipofectamine™ 2000 Transfection Reagent (Invitrogen) following the manufacturer’s instructions.

### Chemicals

The in-house 54-compound library was kindly provided by Professor Neil Carragher (The University of Edinburgh), including FDA-approved drugs and natural products of well-established anti-cancer mechanisms. The concentrations of library compounds were varied from 0.1 to 10 mM according to their optimal effects in previous cell studies in the Carragher’s lab. Ro 08-2750 was purchased from R&D Systems. Oleic acid, quercetin, luteolin, genistein, dihydrotanshinone I (DHTS), CMLD-2, cetylpyridinium chloride (CPC) and gossypol were purchased from Sigma-Aldrich. All 9 reagents were dissolved in DMSO to prepare 20 mM stock solutions.

### Plasmids construction

Human genomic DNA was isolated from HeLa cells using a GenElute™ Mammalian Genomic DNA Miniprep Kit (Sigma-Aldrich). Target genes were PCR amplified from genomic DNA using Phusion^®^ High-Fidelity DNA Polymerase (NEB) and propagated following the instructions of CloneJET PCR Cloning Kit (Thermo). The HuR open reading frame was inserted downstream from the mCherry gene between XhoI and EcoRI in a pJW99 plasmid. DNA segments encoding miRNA stem loop sequences were cloned into a pCG plasmid between the XbaI and BamHI cleavage sites. The 3’-UTR of human α-Syn mRNA gene was inserted into a psiCHECK-2 plasmid between the XhoI and NotI sites downstream the Renilla luciferase (Rluc) gene. Sequence confirmation was carried out in Edinburgh Genomics or GENEWIZ.

### Mutagenesis PCR

Mutants of psiCHECK-2-α-Syn-3’UTR were generated using mutagenesis PCR, where the nucleotides in the 3^rd^ position of the putative miRNA binding sites were mutated. Primers used for mutagenesis are as follows:

miR-7_s1-F: accatcagcagtgattgaagt R: agacttcgagatacactgtaaa
miR-7_s2-F: attaatgatactgtctaagaataatg R: agacacctaaaaatcttataatatat
miR-7_s3-F: acccttaatatttatctgacggta R: agaaacactttaaaggagaatttg
miR-133b_s1-F: gaaacacttaaacaaaaagttcttta R: gtcccaaataaactattaagatatat
miR-153_s1-F: ccacctataaatactaaatatgaaatt R: atagtttcatgctcacatatttttaa
miR-153_s2-F: ccactggttccttaagtggctg R: ataccaaaacacacttctggca
miR-153_s3-F: ccactagtgtgagatgcaaaca R: agagattctgaaaaagacccca

PCR reactions were run for 18 cycles using Phusion polymerase. The template plasmids were digested by DpnI (NEB) treatment at 37°C for 1.5 hour, followed by T4 Polynucleotide Kinase (NEB) treatment at 37°C for 1 hour. Blunt-end ligation was performed using T4 DNA Ligase (NEB) at 4°C overnight. The ligation products were transformed and propagated in *E. coli*.

### Western blot analysis

Cultured cells were harvested and resuspended in Roeder D (200 mg/ml glycerol, 100 mM KCl, 0.2 mM EDTA, 100 mM Tris pH 8.0, 500 μM DTT and 200 μM PMSF). Cell lysis was carried out in a Bioruptor^®^ Plus sonication device (Diagenode) for 10 min (low intensity settings, 30s on/off). 60 μg of proteins in cell lysates were separated on a NuPAGE™ 4-12% Bis-Tris Protein Gel (Invitrogen) and transferred onto a nitrocellulose membrane (GE) in a GENIE^®^ blotter (Idea Scientific) at 12V for 1 h. The membrane was blocked with 1:10 Western Blocking Reagent (Roche) in TBST (20 mM Tris pH 7.5, 137 mM NaCl and 0.1% (v/v) Tween 20). Proteins were detected with the following primary antibodies in TBST containing 1:20 Western Blocking Reagent, including rabbit polyclonal anti-HuR (Millipore), rabbit polyclonal DHX9 antibody (Proteintech), monoclonal anti-α-tubulin antibody (Sigma-Aldrich) and purified mouse anti-α-Synuclein (BD Biosciences). Following three washes in TBST, the membrane was incubated in the horseradish peroxidase (HRP) conjugated secondary anti-rabbit or anti-mouse IgG antibodies (Cell Signalling Technology) and developed with chemiluminescent substrate (Thermo #34580). Quantification of western blot bands were carried out in Image Studio Lite Ver 5.2.

### RNA quantification

Total RNA was isolated from cells or cell extracts following manufacture’s instruction of TRI Reagent™ Solution (Invitrogen) or TRI Reagent^®^ LS (Sigma-Aldrich), respectively. To quantify miRNA levels, reverse transcription and quantitative real-time PCR (qRT-PCR) were performed with miScript II RT Kit and SYBR Green PCR Kit respectively (QIAGEN). To quantify α-Syn mRNA levels, the GoTaq^®^ 1-Step RT-qPCR System (Promega) was used with the primers (F: 5’-gttgtggctgctgctgagaaa; R: 5’-tccctccttggttttggagcctac). The qRT-PCR reactions were performed in a Roche LightCycler^®^ 96 System.

### Dual luciferase assays

Luciferase reporter assays were performed according to the manufacture’s instruction of Dual-Luciferase^®^ Reporter Assay System (Promega). 1.2×10^4^ of HeLa cells were plated in a well of 96-well plates. For each well, 30 ng of psiCHECK2-α-Syn-3’UTR and 16.7 ng of pCG-pri-miRNA plasmids were co-transfected into HeLa cells. Cells were lysed 48 hours after transfection by Passive Lysis Buffer (Promega). The luminescence levels of firefly and Renilla luciferases were recorded by a PolarStar OPTIMA Multidetection Microplate Reader (BMG LABTECH).

### RP-CONA

Ni-NTA agarose beads (Qiagen, 30250) were sieved using 100 μm cell strainers (Falcon, 352360) and 120 μm pore size filters (Millipore, NY2H04700) to obtain a uniform bead size population. The sieved beads were washed with binding buffer (0.3 M NaCl, 20 mM HEPES pH 7.5, 0.01% Triton X-100) and resuspended to 10% slurry. For each reaction, 150 pmol of 6×His-streptavidin (ProteoGenix) was mixed with 5 μl of sieved beads (10%) in a total volume of 20 μl binding buffer at 4°C, 1,000 rpm for 20 min. The biotinylated FITC-pri-miR-7-CTL (5’-FITC-uguuguuuuuagauaacuaaaucgacaacaaa-Biotin-3’) were synthesized by IDT. 40 pmol of RNA was mixed with 5 μl of the streptavidin beads (10%) in a total volume of 20 μl PBS containing 0.01% Triton X-100 at 4°C, 1,000 rpm for 20 min.

HuR KO-HEK293T cells were plated in a p150 dish and transfected with 40 μg of pJW99-HuR plasmids. The cell lysates were obtained 24 hours after transfection by sonication. The concentration of mCherry-HuR in the lysates was quantified following the instruction of an mCherry quantification kit (BioVision). The lysates containing 300 nM of mCherry-HuR were diluted in 20 μl of no glycerol-Roeder D (100 mM KCL, 0.2 mM EDTA, 100 mM Tris pH 8.0, 0.5 mM DTT and 0.2 mM PMSF), and mixed with 30 μl of pulldown solution (1.5 mM MgCl2, 25 mM creatine-phosphate, 0.5 mM ATP, 0.25 μl Ribolock RNase Inhibitor (Invitrogen)) before loaded to a well of a black 384-well plate (SWISSCI or Greiner). 0.5 μl of compounds (dissolved in DMSO at 100 times of the final concentration) were added to the cell lysates and mixed vigorously at room temperature, 1,500 rpm for 20 min.

5 μl of FITC-pri-miR-7-CTL-beads (10%) was added to the treated lysates and mixed at room temperature, 1,500 rpm for 2 hours. Images were acquired in an Opera^®^ High Content Screening System (PerkinElmer) or ImageXpress Micro Confocal (Molecular Devices) at 30 μm above well bottom, 20× magnification with air lenses. On the Opera HCS, three channels were detected: brightfield (690 nm diode at 70% power, 690/70 detection filter, 160 ms exposure time), FITC (488 nm laser at 250 μW, 520/35 detection filter, 120 ms exposure time) and mCherry (561 nm laser at 1250 μW, 600/40 detection filter, 240 ms exposure time). In ImageXpress, two channels were detected: FITC (FITC filter set: excitation 475/34, emission 536/40, exposure 25 ms) and mCherry (Texas Red filter set: excitation 560/32, emission 624/40, exposure 85 ms). FITC and mCherry rings were detected and the ring intensities were quantified in Image J, using a custom plugin (DOI:10.5281/ZENODO.4302193).

### RNA pull-down assay

500 pmol of pri-miR-7-CTL (5’-uguuguuuuuagauaacuaaaucgacaacaaa) was treated with 100 mM NaAc and 5 mM sodium (meta)periodate in 200 μl of water and rotated for 1 hour at room temperature in the dark. The RNA was precipitated by adding 600 μl of 100% ethanol and 15 μl of 3 M NaAc in dry ice for 20 min, followed by centrifugation at 13,000 rpm, 4°C for 10 min. The RNA pellet was washed with 70% ethanol and resuspended in 500 μl of 100 mM NaAc pH5. 200 μl of adipic acid dihidrazide-agarose (Sigma) were washed with 100 mM NaAc and mixed with 500 μl of the periodate oxidized pri-miR-7-CTL overnight at 4°C in the dark. The pri-miR-7-CTL-beads were washed by 2M KCL, Buffer G (20 mM Tris-HCl pH 7.5, 137 mM NaCl, 1 mM EDTA, 1% Triton X-100, 10% glycerol, 1.5 mM MgCl2, 1 mM DTT, and 200 μM PMSF) and Roeder D respectively.

250 μl of HeLa cell lysates containing 1 mg of total protein were pre-incubated with 100 μM of test compounds and 400 μl of pulldown solution at 37°C, 700 rpm for 20 min. The pri-miR-7-CTL-beads were incubated with the treated cell lysates, at 37°C, 1,200 rpm for 30 min. After washing with Buffer G, the beads were mixed with 6 μl of NuPAGE™ Sample Reducing Agent, 15 μl of LDS Sample Buffer and 39 μl of water. Proteins captured by RNA were denatured at 70°C, 1,000 rpm for 10 min. 30 μl of the supernatant was loaded onto an SDS-PAGE gel and western blot was performed to detect the level of proteins.

### Generation of gene knockout cells

To knock out HuR in HEK293T and HeLa cells, a pair of guide RNAs (5’-cgaagucuguucagcagcau and 5’-cuugggucauguucugaggg) targeting the exon2 of human HuR gene were designed. The Alt-R^®^ CRISPR-Cas9 crRNAs and tracRNA were synthesized by IDT. 100 μM of each crRNA was mixed with 100 μM of tracRNA in 100 μl duplex buffer (IDT) at 95°C for 5 min, to form two crRNA-tracRNA duplexes. HEK293T or HeLa cells were seeded in a 24-well plate. 1.5 μl of each duplex were co-transfected with 1 μl of GeneArt™ CRISPR Nuclease mRNA (Invitrogen). The cells were diluted and aliquoted to 96-well plates to make less than 1 cell count per well and lysed by 30 μl of Passive Lysis Buffer (Promega) when confluent. 2 μl of the cell extracts were loaded onto the nitrocellulose membranes and tested against HuR and DHX9 antibodies. Cells showing HuR negative and DHX9 positive were selected and confirmed using western blot analysis. The genomic DNA of the knockouts were extracted and fragments covering the expected HuR knockout sites were PCR amplified using the forward primer (5’-gccctggacagtacactcgcc) and reverse primer (5’-ccacatggccgaagactgca). After ligating into the cloning vector using the CloneJET PCR Cloning Kit (Thermo), the DNA fragments were sequenced, and the mutations were identified.

To knock out miR-7 in HeLa cells, a pair of guide RNAs (5’-acauucaauacuaaucuugc and 5’-accaaucauuuguccuguag) were designed flanking the stem loop sequence of human pri-miR-7-1 gene. Transfection of the CRISPR-Cas9 system was carried out as described before. Crude DNA was extracted by dissolving cells in solution 1 (25 mM NaOH, 0.2 mM EDTA) at 98°C for 1 hour, and terminated by equal volume of solution 2 (40 mM Tris-HCL pH5.5). PCR was performed with primers flanking the targeted region (F: 5’-ctgcagaacaggtcagtttaagtt, R: 5’-tgcagaacacctatgaagcaga). The PCR products were visualised on an agarose gel and cells generating band shifts were selected. The miR-7 levels were tested by qRT-PCR. Sequences were determined using PCR products amplified from purified genomic DNAs of putative miR-7 knockouts.

## RESULTS

### MiR-7 is a major inhibitor of α-Syn expression

MiR-7 and other miRs, such as miR-133b, miR-153, miR-34b or miR-34c, which also have the potential to target a-Syn, have been shown to be downregulated in PD, thus allowing a-Syn overproduction and accumulation (23,25,43–45). We collated all validated and predicted miRs targeting the 3’-UTR of α-Syn mRNA and cloned their pri-miR sequences into the pCG expression vector (Figure 1A). We used a dual luciferase reporter assay in HeLa cells with overexpression of individual miRs targeting α-Syn mRNA 3’-UTR coupled with Renilla luciferase mRNA. This assay showed that only miR-7 exhibited significant inhibition (>50%) of Renilla luciferase compared to other miRs, when equal amounts of pri-miR plasmids were transfected (Figure 1B). The upregulation of mature miR-7 was equal or less when compared with other miRs (Supplementary Figure S1A and data not shown). Moreover, a single-nucleotide mutation in the previously identified miR-7 binding site (miR-7_s1) can desensitise α-Syn mRNA 3’-UTR to miR-7 overexpression, while other miR-7 binding sites (miR-7_s2 and miR-7_s3) did not present significant differences between the wildtype and mutated reporters (Figure 1C). Interestingly, mutations of miR-133b_s1 and miR-153_s1 sites exerted significant upregulation of expression from the α-Syn mRNA 3’-UTR Renilla luciferase construct, proving that miR-133b and miR-153 are involved in regulating a-Syn levels, albeit to lesser extent and with more complex regulatory networks (Figure 1C).

**Figure 1.**
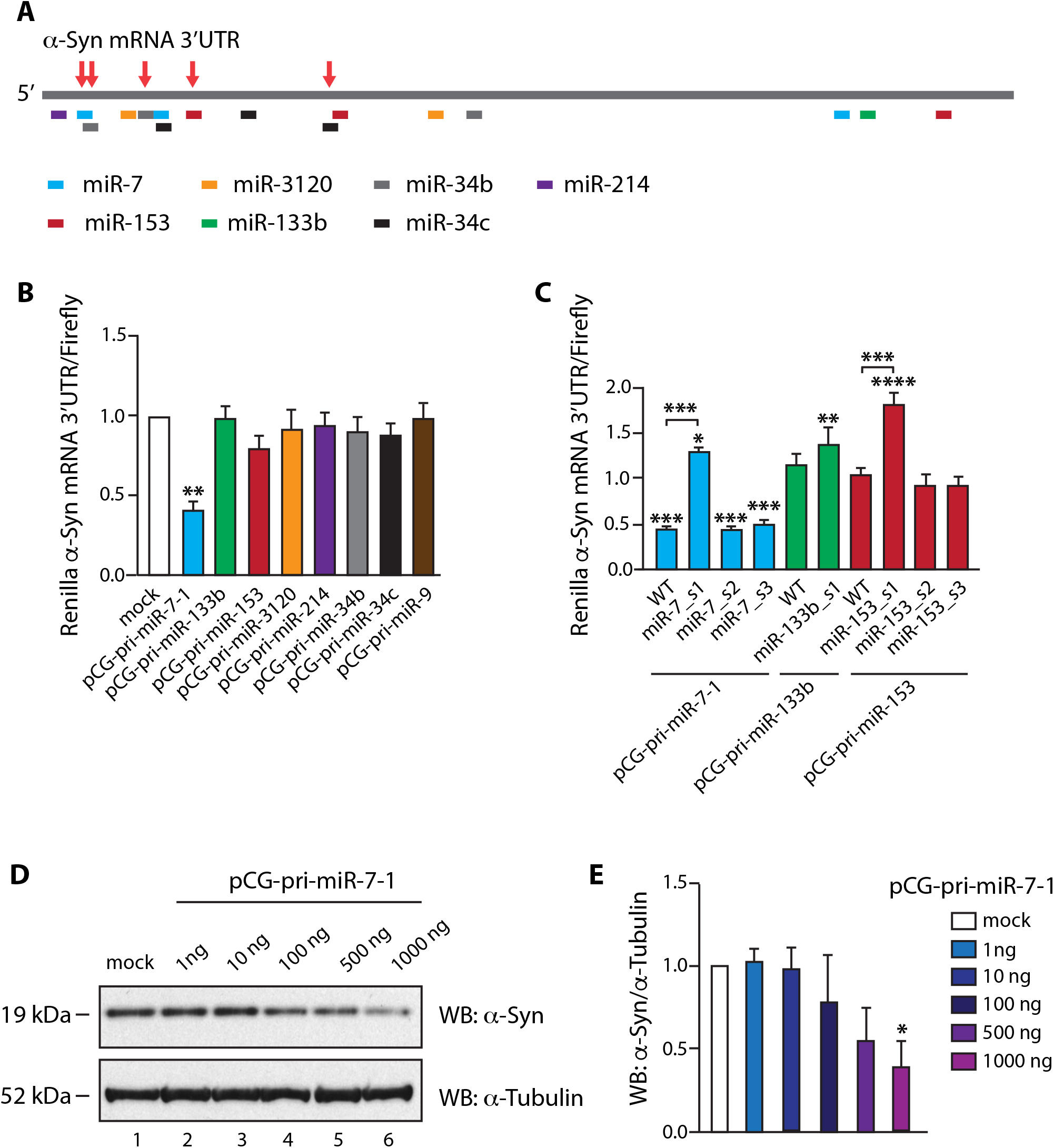
MiR-7 is a major suppressor of α-Syn expression. (**A**). Predicted binding sites of miRs on the 3’-UTR of α-Syn mRNA (~2,500 bp). The putative binding sites of different miRs are annotated at the approximate positions on the α-Syn mRNA 3’-UTR. These sites were predicted by TargetScan (44) or provided by miRTarBase (45). Sites highlighted by the red arrows were previously reported. (**B**). MiR-7 inhibited luminescence of a dual luciferase reporter bearing the gene of α-Syn mRNA 3’-UTR downstream the Renilla gene. Equal amount of each pCG-pri-miR plasmid was co-transfected with the luciferase reporter. PCG-pri-miR-9 was tested as a negative control. Luciferase levels were recorded 48 hours after transfection. Mean Renilla/firefly values and SEM from 6 independent repeats are shown. (**C**). Single-nucleotide mutation of miR-7 binding site inactivated the inhibitive effects of miR-7. The nucleotide on the 3^rd^ position of miR-7, miR-153 or miR-133b targeted seed region on α-Syn mRNA 3’-UTR gene was mutated individually. The mutants were numbered according to the binding sites from 5’ to 3’ of α-Syn mRNA 3’-UTR. Luciferase levels were measured 48 hours after the co-transfection of pCG-pri-miR-7-1, pCG-pri-miR-153 or pCG-pri-miR-133b plasmids with reporters bearing wildtype or mutated α-Syn mRNA 3’-UTR gene. The luciferase levels were relative to co-transfection of pCG-pri-miR-9 with wildtype reporter. Mean Renilla/firefly values and SEM from three independent repeats are shown. (**D, E**). α-Syn expression was inhibited by upregulated miR-7. 1: Mock HeLa cells without DNA transfected. 2-6: An increasing amount of pCG-pri-miR-7-1 was transfected into HeLa cells. The expression of α-Syn and α-tubulin were detected by western blot 48 hours after transfection. Relative α-Syn/α-tubulin levels were normalised to mock. Mean values and SEM from three independent repeats are shown. Statistically significant differences compared to mock were interpreted by SPSS one-way ANOVA, *P<0.05, **P<0.01, ***P<0.001, ****P<0.0001.

Importantly, a gradient of miR-7 overexpression exerted gradual and significant reduction of the expression of endogenous α-Syn in HeLa cells (Figure 1D, E), which is consistent with a previous publication (23). HeLa cells were chosen because according to Human Protein Atlas these cells have high basal levels of α-Syn transcripts when compared with most other cultured cells (46). Other miRs, exemplified by miR-153 and miR-133b, exerted no significant inhibitive effects on endogenous α-Syn levels, in spite of their previously reported function as α-Syn inhibitors (Supplementary Figure S1B) (47,48). In summary, we conclude that miR-7 is the most potent α-Syn suppressor and uses a well-defined target site on the α-Syn mRNA 3’-UTR.

### RP-CONA: a novel, lysate-based scanning microscopy on-bead screening platform for small molecules that modulate RNA-protein complexes

Due to the supremacy of miR-7 in inhibiting the expression of α-Syn, it will be crucial to pursue therapeutic approaches for PD through the miR-7/α-Syn pathway. We hypothesize that as the biogenesis of pri-miR-7-1 is inhibited by HuR through an interaction with the conserved terminal loop (CTL), small molecules that dissociate HuR from miR-7 CTL will facilitate the production of mature miR-7, which in turn will contribute to a reduction of α-Syn levels (32).

To identify pri-miR-7/HuR inhibitors, we developed a screening platform that combines techniques of RNA pull-down (RP) from eukaryotic cell extracts, and on-bead scanning confocal microscopy (CONA), given the name RP-CONA (Figure 2). First, we attached 5’FITC and 3’biotin-tagged pri-miR-7-1-CTL to 6×His-Streptavidin Ni-NTA agarose beads. Then we used HuR KO HEK293T cells, generated with CRISPR-Cas9 targeting, to overexpress mCherry-HuR (Supplementary Figure 2A-C). The HuR KO cells were used to avoid signal dilution from an untagged, endogenous HuR. Next, we performed RNA pull-down and imaged the fluorescently labelled RNA and mCherry-HuR on a confocal imaging system. Fluorescent rings/halos on the periphery of microbeads indicate binding events in separate detection channels. The amounts of bound RNA or protein were detected from fluorescence emission intensities of the rings/halos. Pri-miR-7/HuR inhibitors are expected to attenuate mCherry ring intensities without affecting FITC levels.

**Figure 2.**
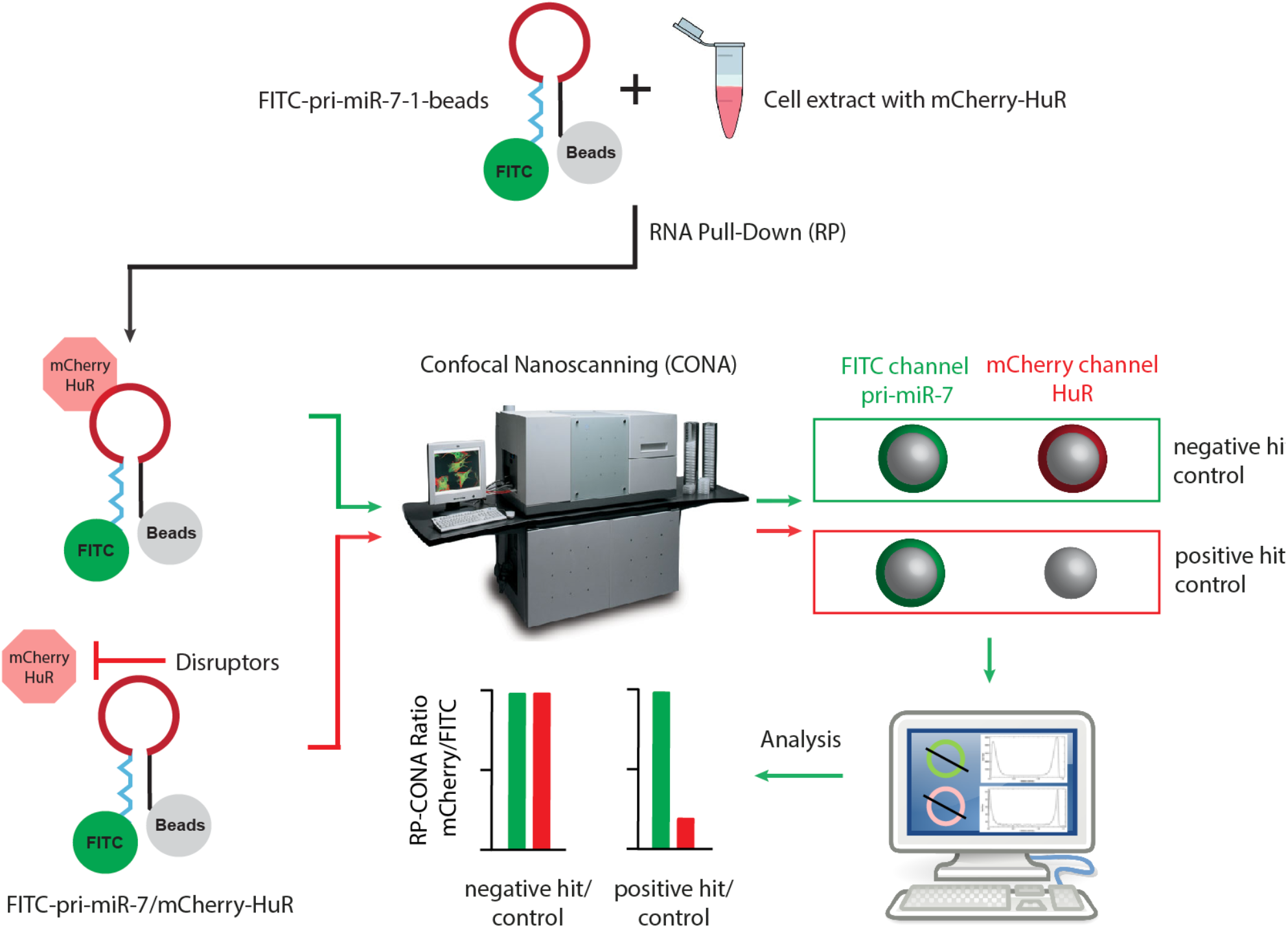
The working scheme of RP-CONA. RNA Pull-down (RP): 5’-FITC-pri-miR-7-1-CTL-biotin-3’ is coupled to streptavidin coated agarose beads. Cell lysates are extracted from HuR KO HEK293T cell overexpressing mCherry-HuR and treated with small molecules. The RNA-coupled beads are incubated with cell lysates to pull-down mCherry-HuR. Confocal Nanoscanning (CONA): Beads are imaged using a confocal image scanning platform. On-bead FITC-pri-miR-7-CTL and mCherry-HuR are detected with high sensitivity as fluorescent rings/halos on the outer shell of bead in corresponding detection channels. Inhibitors are able to attenuate the mCherry fluorescence without affecting FITC signals. Analysis: The fluorescent rings are detected in imageJ. Three measurements are taken across each ring and generate the fluorescent intensity profiles to obtain arbitrary intensity values.

We first tested the RP-CONA assay using the Opera HCS instrument. The preliminary results showed that RP-CONA generated FITC rings in the presence of the fluorescently tagged RNA, and mCherry rings only when mCherry-HuR was pulled down by pri-miR-7-CTL (Figure 3A). No overlapping signals were observed in different detection channels. The addition of an increasing concentration of untagged pri-miR-7-CTL competitively decreased mCherry signals (Figure 3B, C). Oleic acid, the known pri-miR-7/HuR complex inhibitor reduced mCherry ring intensities at high millimolar concentrations, as reported before (Figure 3D, E) (35). To show that RP-CONA can be run on various scanning instruments we then switched to another confocal image platform, the ImageXpress, which gave similar quality of images compared to Opera HCS (Supplementary Figure 3A). Moreover, the relative mCherry/FITC value was proportional to the concentration of mCherry-HuR (Supplementary Figure 3B). These results demonstrate that RP-CONA is a robust technique to screen for RNA-protein interaction modulators and that it could be used with various confocal image scanning platforms.

**Figure 3.**
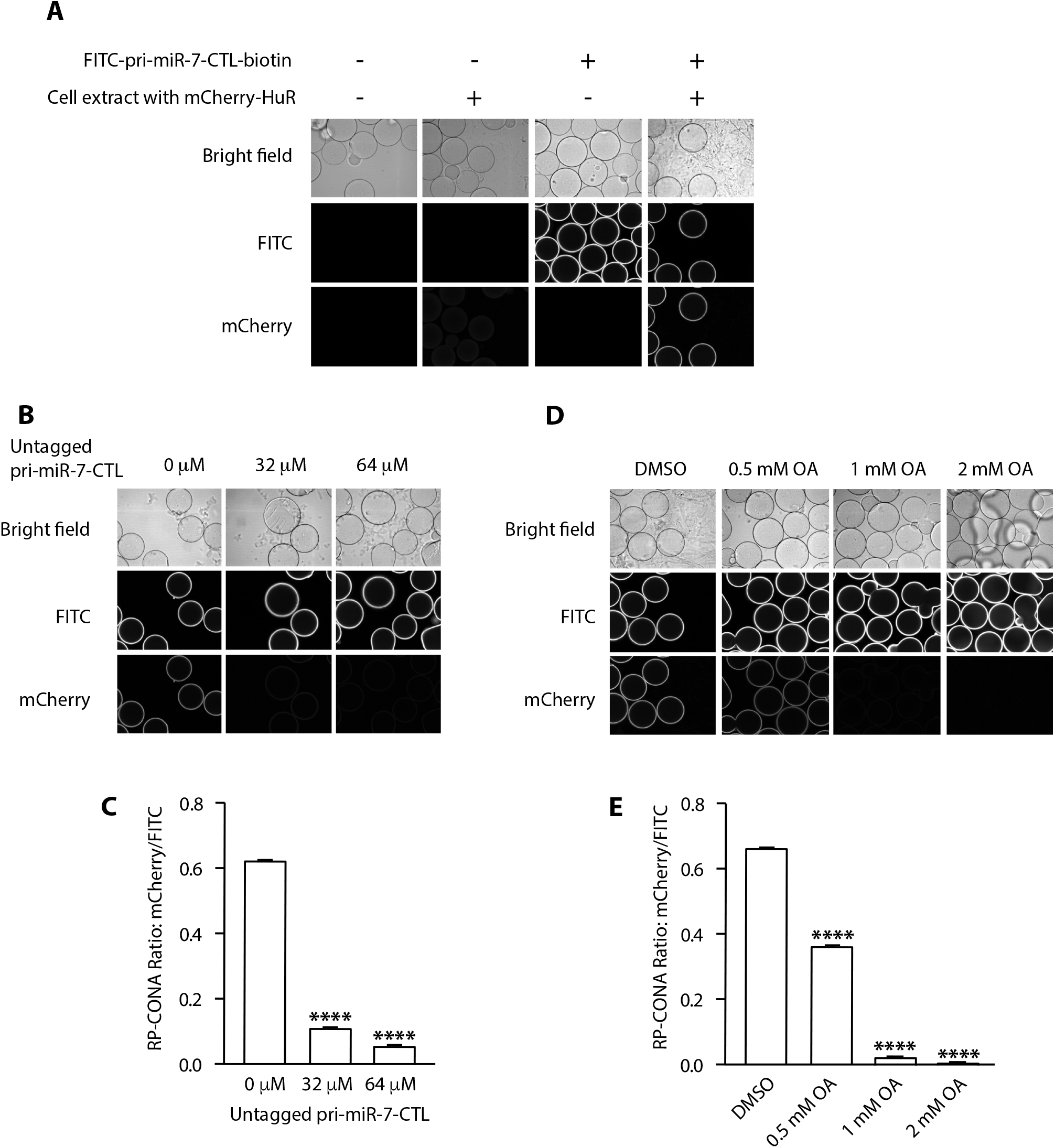
Optimisation of RP-CONA method in the Opera HCS. (**A**). Beads images of blank beads; blank beads incubated with cell lysates containing mCherry-HuR; FITC-pri-miR-7-CTL-beads incubated in lysates-free buffer; and FITC-pri-miR-7-CTL-beads after mCherry-HuR pulldown. (**B, C**). mCherry/FITC signals were reduced by untagged pri-miR-7-CTL. Cell lysates containing mCherry-HuR were treated with an increased concentration of untagged pri-miR-7-CTL before pulldown. (**D, E**). mCherry/FITC signals were reduced by a pri-miR-7/HuR inhibitor. Cell lysates containing mCherry-HuR were treated with DMSO, or an increased concentration of OA before pulldown. All values were obtained from three technical repeats. Mean mCherry/FITC ring intensities and SD between triplicates are shown. Statistically significant differences compared to 0 μM untagged pri-miR-7 or DMSO were interpreted by SPSS independent sample t-test, **** P<0.0001.

### Pri-miR-7/HuR inhibitors identified by RP-CONA screening of a focused library

MSI2 is an essential RBP that assists HuR during the inhibition of miR-7 biogenesis (32). A range of compounds have been recognised to inhibit HuR or MSI2 binding to their target mRNAs (6). By collecting the commercially available HuR/MSI2 inhibitors, we built up a focused library and tested them using RP-CONA at 100 μM concentration (Figure 4A). The Z’ factor of the screen is 0.93, and the coefficient of variations (CVs) of the negative control (DMSO) and positive control (untagged pri-miR-7-CTL) equal to 1.75% and 0.96%, respectively (Figure 4B). As reported previously (35) OA did not show significant disruptive effects on pri-miR-7/HuR complex at 100 μM. Importantly, we have identified quercetin, luteolin and gossypol as pri-miR-7/HuR inhibitors, which generated more than 50% inhibition at 100 μM concentration according to the relative mCherry/FITC signals compared to the DMSO control (Figure 4C). In summary, RP-CONA identified new potential inhibitors of RNA/HuR complexes.

**Figure 4.**
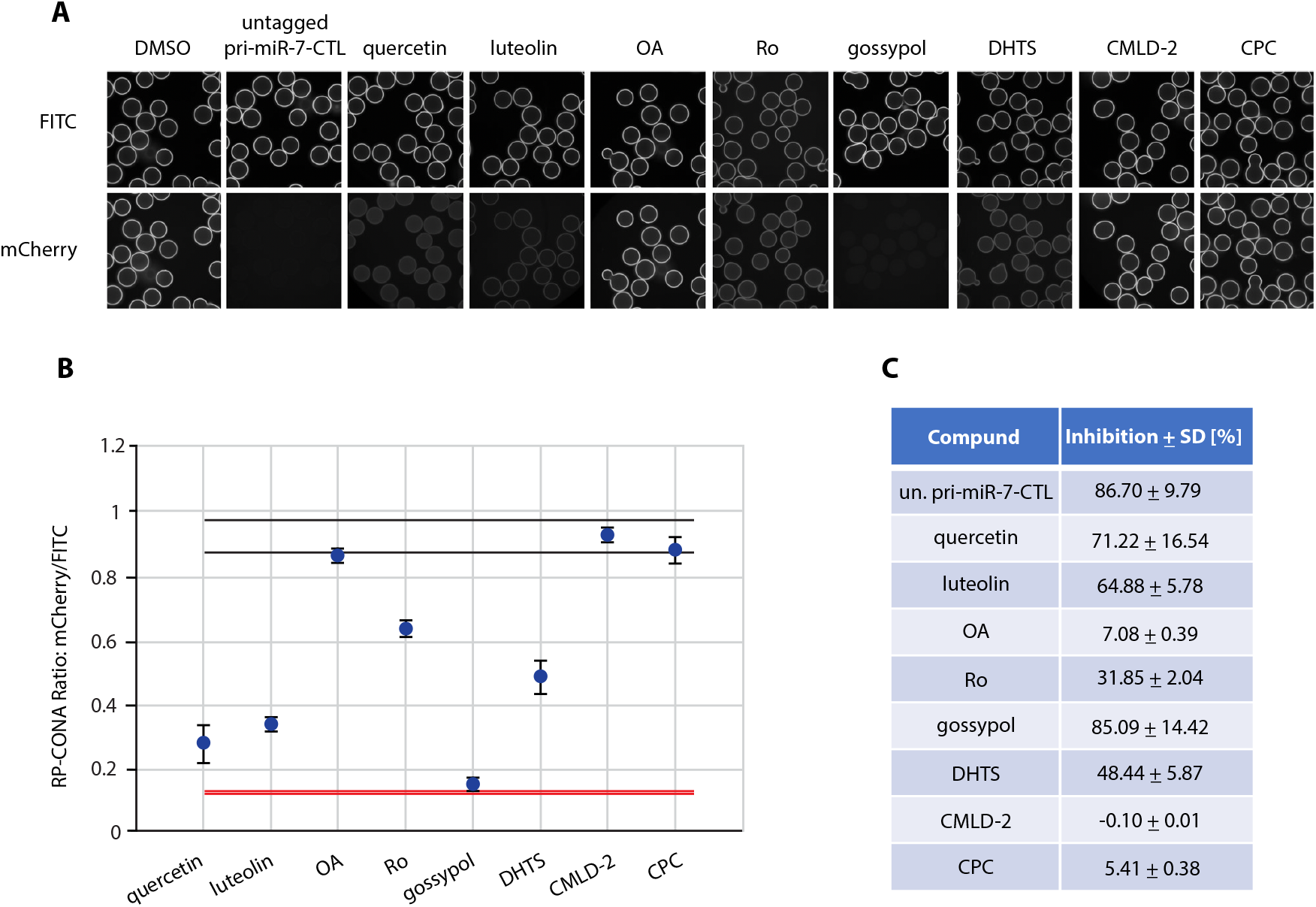
RP-CONA identified pri-miR-7/HuR inhibitors from a focused library. (**A**) Cell lysates containing mCherry-HuR were treated with DMSO, 50 μM of untagged pri-miR-7-CTL or 100 μM of compounds before pulldown. Beads images taken in ImageXpress. (**B**) Relative mCherry/FITC ring intensity mean and SD between the beads in each well after compound treatment are shown. DMSO (RP-CONA ratio: 0.932±0.016, CV: 1.75%, n=5) served as a negative control while 50 μM of untagged pri-miR-7-CTL (RP-CONA ratio: 0.124±0.001, CV: 0.96%, n=5) served as a positive control. Z’ = 0.93. Black lines: DMSO mean ± 3×SD between 5 repeated wells. Red lines: untagged pri-miR-7-CTL mean ± 3×SD between 5 repeated wells. At least 400 beads were included in each control analysis. (**C**). Inhibition of RP-CONA signals compared to DMSO. The percentage inhibition of compounds relative to DMSO mean are shown. The SD of relative inhibitions between the beads in each well after compound treatment are shown.

### A small-scale prototype RP-CONA screen of pri-miR-7/HuR inhibitors

In order to analyse RP-CONA’s ability to scale up and to establish the sensitivity of the method, we performed a small-scale prototype screen using an in-house library containing 54 FDA-approved drugs or natural products, together with 8 compounds from the previous focused screen (Figure 5A). Here, we applied quercetin as a positive control and DMSO as a negative control. The CVs of the negative and positive controls were 6.70% and 2.18%, respectively. Most compounds did not show significant stabilisation or destabilisation of the pri-miR-7/HuR complex. Additionally, we identified genistein as a new hit, albeit not as effective as quercetin or luteolin. To validate the active compounds identified in the primary screen, we tested them in a standard pull-down assay where pri-miR-7-CTL was covalently linked to the beads (49). This assay, unlike RP-CONA, only gives semi-quantitative readout. The western blot analysis of the HuR pull-down confirmed the strong disruptive effects of quercetin and luteolin (Figure 5B). Similar effects from DHTS, Ro, and genistein were also observed. Meanwhile, gossypol largely reduced pull-down of the RNA helicase DHX9, implying the effects of this compound are unspecific. Our most effective inhibitors quercetin and luteolin are close analogues (Figure 5C). We tested quercetin and luteolin in RP-CONA at a range of concentrations. The RP-CONA signals were reduced in a dose-dependent manner for both quercetin and luteolin, showing the half maximal inhibitory concentrations (IC50s) at 2.15±0.16 μM and 2.03±0.25 μM, respectively (Figure 5D). These observations indicate that RP-CONA primary hits were successfully validated using a classic biochemical method and further confirm quercetin and luteolin as the most promising inhibitors of the pri-miR-7/HuR complex.

**Figure 5.**
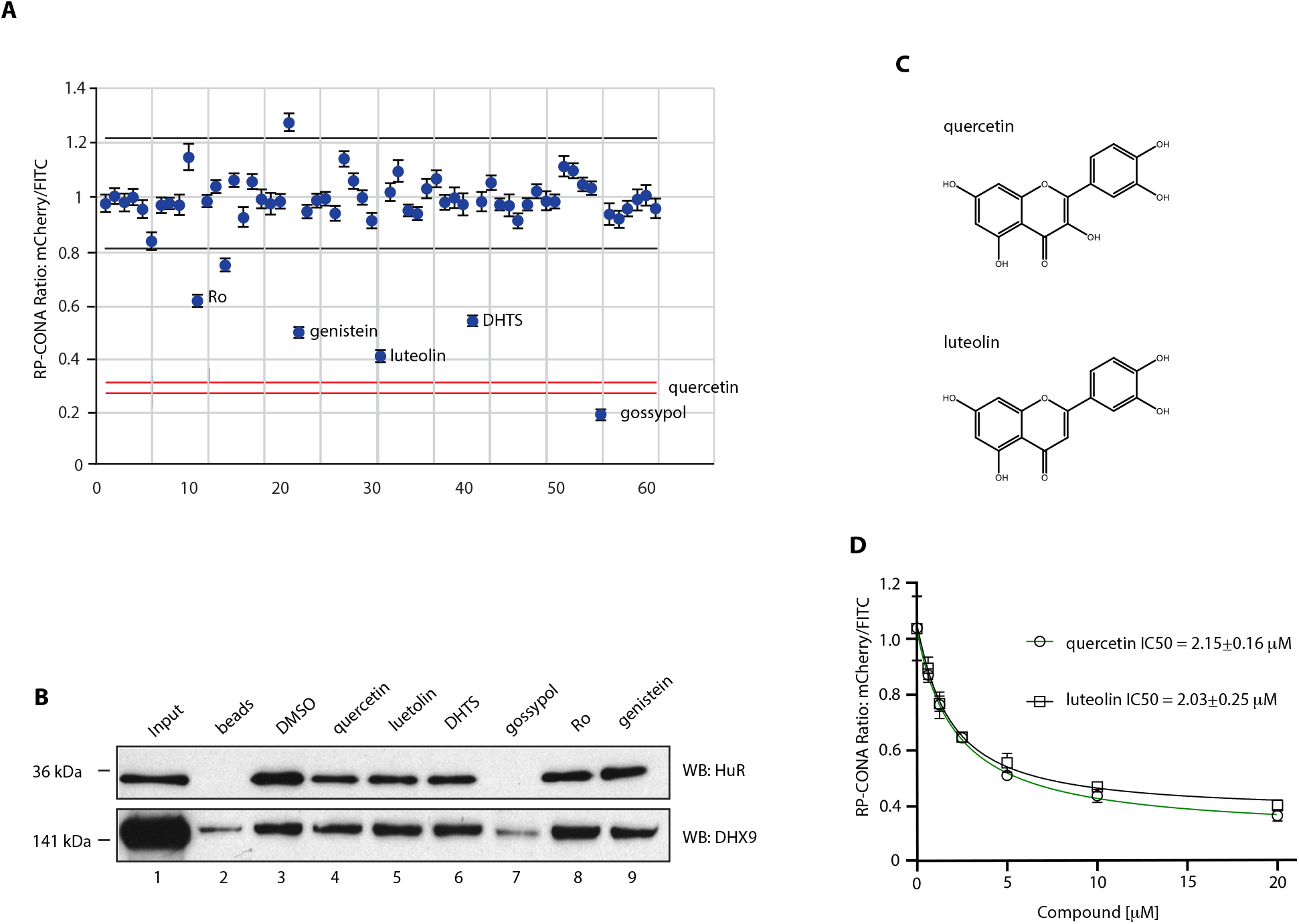
Identification and validation of pri-miR-7/HuR inhibitors from an enlarged library. (**A**) A small-scale prototype RP-CONA screen to test the disruptive effects of 61 compounds (54 from an in-house library varied from 1 to 100 μM, and 7 from the previous identified HuR or MSI2 inhibitors at 100 μM) on pri-miR-7/HuR. Relative mCherry/FITC ring intensity mean and SD between the beads in each well after compound treatment are shown. DMSO (RP-CONA ratio: 1.009±0.067, CV: 6.70%, n=6) served as a negative control while 100 μM of quercetin (RP-CONA ratio: 0.288 ±0.006, CV: 2.18%, n=6) served as a positive control. Z’ = 0.69. Black lines: DMSO mean ± 3×SD between 6 repeated wells. Red lines: quercetin mean ± 3×SD between 6 repeated wells. At least 500 beads were included in each control analysis. Compounds generating larger than 40% (6 times CV of negative controls) inhibition are annotated. (**B**) Disruption of endogenous HuR pulled down by pri-miR-7-CTL. Pri-miR-7-CTL was covalently linked to agarose beads. HeLa extracts were treated with DMSO or 100 μM of each compound prior to pull-down. HuR and DHX9 in the pull-down components were detected by western blot. 1: Input. Unbound proteins in the lysates after pulldown. 2: Proteins pulled down by beads without RNA. 3-9: Proteins pulled down by pri-miR-7-CTL-beads after compound treatment. (**C**) Chemical structures of quercetin and luteolin. (**D**) Quercetin and luteolin disrupted pri-miR-7/HuR in a dose-dependent manner in RP-CONA. Increased concentrations of quercetin and luteolin were tested in RP-CONA. Relative mCherry/FITC ring intensity mean and SD between triplicated wells after compound treatment are shown. The RP-CONA signals were curve fitted by non-linear regression-four parameter [inhibitor] versus response, and IC50s were determined with the 4-parameter equation in GraFit v7.0.3 (Erithacus Software Limited) (41). The IC50s of quercetin and luteolin were 2.15±0.16 μM and 2.03±0.25 μM, respectively.

### Quercetin inhibits cellular α-Syn and upregulates miR-7

Finally, we tested quercetin and luteolin in HeLa cells at 20 μM concentration. Quercetin treated cells had a significantly reduced level of α-Syn protein, with a 1.5-fold upregulation of mature miR-7 level (Figure 6A-C). Interestingly, luteolin had no significant effect on the α-Syn or miR-7, suggesting different bioavailability, dynamics or cellular metabolism when compared with quercetin. To find out if quercetin acts through the miR-7/α-Syn pathway, we deleted the stem loop region of pri-miR-7-1 in the genome of HeLa cells, using CRISPR-Cas9 and generated miR-7^+/-^ and miR-7^-/-^ cell lines (Supplementary Figure S4). However, quercetin could still inhibit the expression of α-Syn, even though miR-7 was reduced by half or completely depleted (Figure 6D). Next, we tested these compounds in HuR KO HeLa cells generated by CRISPR-Cas9 (Supplementary Figure S2 A, B, D). Importantly, in these cells, quercetin did not affect the levels of α-Syn expression (Figure 6E, F). Intriguingly, in HeLa HuR KO cells the miR-7 levels were upregulated by 2 and 3-fold by quercetin and luteolin, respectively (Figure 6G). This suggests more complex network of miRNA regulation in the absence of HuR. Finally, we investigated the mRNA levels of α-Syn after quercetin treatment. Quercetin significantly reduced α-Syn mRNA by 50% in wildtype HeLa and miR-7 KO cells, while a less significant 20% inhibition was seen in HuR KO HeLa cells (Figure 6H). Thus, we conclude that the HuR-mediated activity of quercetin towards α-Syn is exerted at mRNA level. Further research will show how quercetin regulates α-Syn mRNA abundance. In summary, we conclude that in our cellular model quercetin downregulates α-Syn expression and this regulation is HuR-dependent.

**Figure 6.**
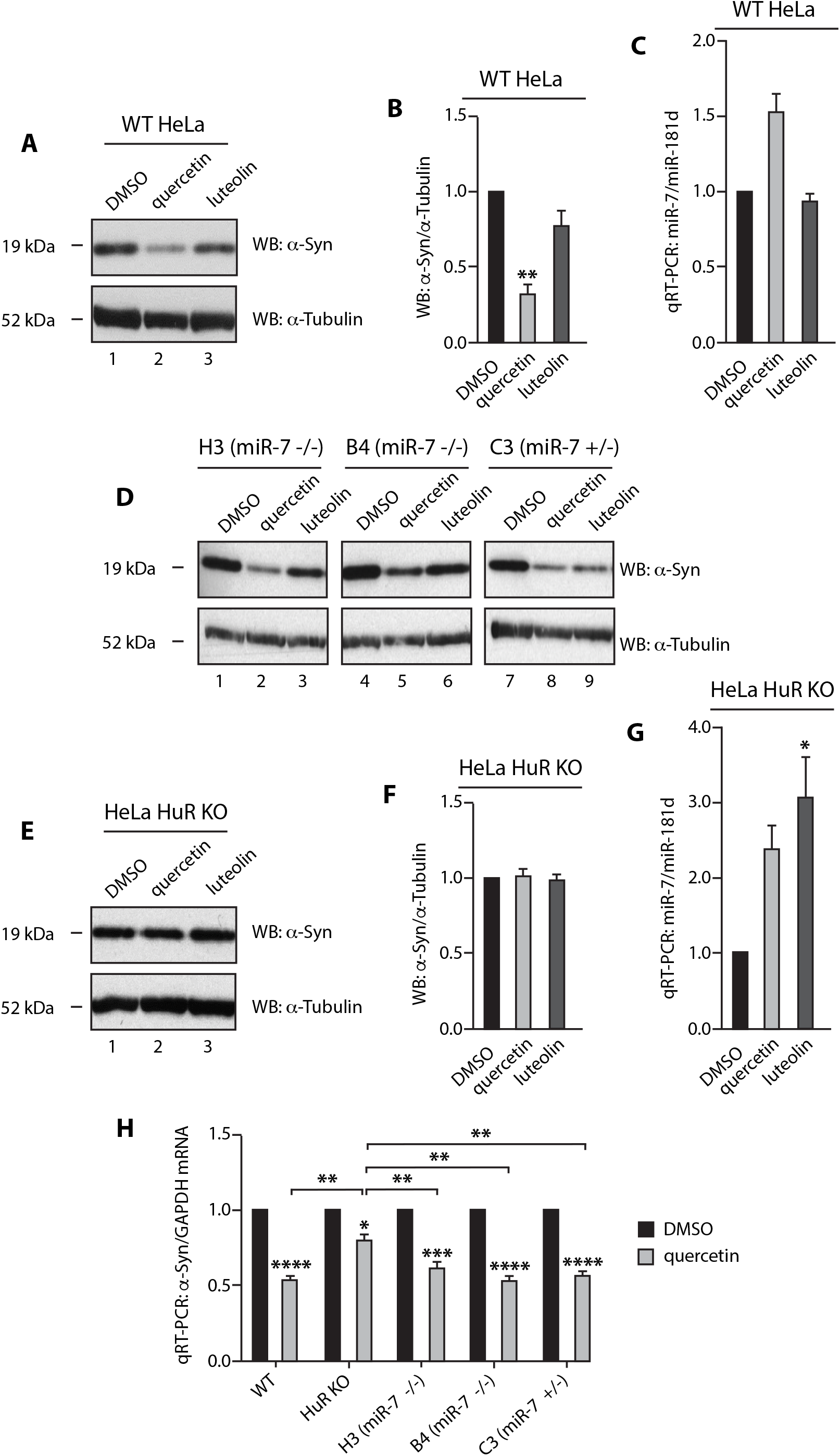
Quercetin inhibits cellular α-Syn and upregulates miR-7. (**A, B**). Quercetin significantly downregulates α-Syn expression in HeLa cells. HeLa cells were treated with 1: DMSO, 2: 20 μM of quercetin or 3: luteolin and harvested 48 hours after treatment. Levels of α-Syn and α-tubulin were detected by western blot. Mean α-Syn/α-tubulin and SEM from three independent repeats are shown. (**C**). Quercetin upregulates cellular miR-7 level by 1.5-fold. HeLa cells were treated with DMSO, 20 μM of quercetin or luteolin and harvested 48 hours after treatment. Mature miR-7 and miR-181d (housekeeping miR control) levels were determined by qRT-PCR. Mean miR-7/miR-181d and SEM from three independent repeats are shown. (**D**) Quercetin reduces α-Syn expression in a miR-7-independent pathway. HeLa miR-7 KO H3 (miR-7^-/-^), B4 (miR-7^-/-^) and C3 (miR-7^+/-^) were treated with 1, 4, 7: DMSO, 2, 5, 8: 20 μM of quercetin, or 3, 6, 9: luteolin for 48 hours. Expression of α-Syn and α-tubulin were detected by western blot. (**E, F**) Quercetin reduces α-Syn expression in an HuR-dependent pathway. HuR KO HeLa cells were treated with 1: DMSO, 2: 20 μM of quercetin or 3: luteolin and harvested 48 hours after treatment. Levels of α-Syn and α-tubulin were detected by western blot. Mean α-Syn/α-tubulin and SEM from three independent repeats are shown. (**G**) Quercetin and luteolin upregulates cellular miR-7 level in HuR KO HeLa. HuR KO HeLa cells were treated with DMSO, 20 μM of quercetin or luteolin and harvested 48 hours after treatment. Mature miR-7 and miR-181d levels were determined by qRT-PCR. Mean miR-7/miR-181d and SEM from three independent repeats are shown. (**H**) Quercetin downregulates α-Syn mRNA level in an HuR-dependent manner. Wildtype, HuR KO and miR-7 KO (H3, B4 and C3) HeLa were treated with DMSO or 20 μM of quercetin and harvested 48 hours after treatment. α-Syn and GAPDH mRNA levels were determined by qRT-PCR. Mean α-Syn/GAPDH and SEM from three independent repeats are shown. Statistically significant differences compared to DMSO, or between quercetin treated cells were interpreted by SPSS independent sample t-test, * P<0.05, ** P<0.01, *** P<0.001, **** P<0.0001.

## DISCUSSION

α-Syn has become a popular target for investigational PD therapies, with RNA interference (RNAi) strategies focusing on α-Syn repression presently at preclinical stages (50). An shRNA therapeutic achieved a 35% knockdown of SNpc α-Syn, and showed protective effects in a PD rat model without notable toxicity (51). However, neuronal toxicity has been found in animal brains when another siRNA-induced α-Syn knockdown reached 90% (52,53). Therefore, the knockdown efficiency of α-Syn-targeted RNAi needs to be tightly controlled within an appropriate range in order to reach a compromise of both safety and efficacy (53,54). MicroRNAs, known as fine tuners of gene expression, buffer the expression network against environmental or genetic stress (55). Here we provide evidence that miR-7 is the most effective suppressor of α-Syn expression, among all potential α-Syn targeting miRs that are downregulated in PD (Figure 1). Therefore, fine-tuning α-Syn expression by elevating miR-7 level could provide a novel avenue for the development of PD therapy.

Previously we have shown that miR-7 biogenesis is inhibited through an interaction between the pri-miR-7-1-CTL and HuR (32). The best elucidated miR post-transcriptional regulation is between let-7 and Lin28 proteins. A range of screens targeting let-7/Lin28 have been carried out using fluorescence resonance energy transfer (FRET) or fluorescence polarisation (FP) methods, and some hits have shown promising anti-cancer properties by dissociating the RNA-protein interaction and upregulating let-7 levels (6). However, due to a lack of understanding of the interaction domains where pri-miR-7 binds HuR, it seems challenging to identify pri-miR-7/HuR inhibitors using similar approaches. To tackle this problem, we developed the RP-CONA technique that combines the RNA pull-down assay in human cell extracts with the CONA screening platform (38–40).

The RNA pull-down assay captures proteins by on-bead RNA fragments from eukaryotic cell lysates (38). The application of cell lysates instead of purified protein allows the screening to take place in a more physiological environment and might help avoiding false positives that are not functional *in vivo*. Moreover, this technique removes the need of protein purification, making it an alternative solution to study RNA/protein binding events for those proteins with difficulty in purification. CONA is a previously established, sensitive and quantifiable on-bead imaging platform (56,57). The only known ligand targeting the RNA recognition motif-3 (RRM3) of HuR was identified by CONA from one-bead-one-compound libraries (58). Notably, using the RRM3 binder and CONA, an ATP-binding pocket was discovered in RRM3, which was not detected using conventional binding assays (58,59). CONA is also highly flexible, and it can be modified to monitor complex biological processes in real time, such as ubiquitination-related enzymatic activities and aggregation of α-Syn (39,41). A recent publication successfully identified protein binders from natural product extracts with CONA (37). Here, for the first time we applied CONA to study RNA-protein interactions by using a lysate-based screening strategy. We believe that this combination has a high potential for further exploitation in miniaturized drug screening.

In order to look for pri-miR-7/HuR inhibitors, we started with a focused prototype screen. The library contained 8 compounds that were known to interfere with the RNA binding activities of HuR or MSI2, as both of the proteins are essential during the blockage of miR-7 biogenesis (6). We found half of the compounds resulting in more than 50% inhibition of RP-CONA signals (Figure 4). By integrating the focused library with a diverse in-house 54-compound library, we have shown that the former 4 compounds are still effective, and only one additional hit was discovered from the new library. The disruptive effects of these hits were confirmed in RP-CONA with a gradient of concentrations, and the IC50s of the most effective hits quercetin and luteolin were around 2 μM. We subsequently validated the primary hits using the standard RNA pull-down assay followed by western blot, and managed to exclude the unspecific disruptor gossypol, which also interrupted the binding of the reference protein DHX9 (Figure 5).

Quercetin has been shown to inhibit the interaction between HuR and TNF-α mRNA, with an IC50 of 1.4 μM in the RNA electrophoretic mobility gel shift assay (EMSA) (42). Luteolin binds the RRM1 of MSI1, the paralogue of MSI2, with a dissociation constant (Kd) of around 3 μM (60). Interestingly, luteolin is a close analogue of quercetin. In an FP assay, both luteolin and quercetin interrupted MSI1-RRM1 binding with a short RNA motif at micromolar and millimolar IC50s, respectively (60). These findings imply that quercetin is likely to be a dual-inhibitor of both HuR and MSI2, marking it a strong candidate as an miR-7 enhancer. The IC50 value derived from RP-CONA for quercetin is 2.15 μM (Figure 5D), which closely matches the value from previous EMSA analysis. Importantly, we identified a 1.5-fold upregulation of mature miR-7 in HeLa cells after 20 μM of quercetin treatment (Figure 6C). This is consistent with what we saw when HuR or MSI2 or both were knocked down (32). At the same time, a significant downregulation of α-Syn expression was observed (Figure 6A, B). It has been reported that quercetin exerts neuroprotective effects in different cell or animal models of PD (61–63). Moreover, oxidized quercetin can prevent α-Syn fibrillization *in vitro* (64). Interestingly, in neuron-like PC12 cells α-Syn expression was induced by 50 to 500 μM of quercetin, albeit reduced when quercetin reached 1 mM (65). These discrepancies may be attributed to the selection of cell lines. We also used quercetin to treat human neuroblastoma SH-SY5Y and neuronlike mouse embryonic carcinoma P19 cell lines. The reduction of α-Syn expression was notable but was less significant than that observed in HeLa cells (data not shown). As mentioned above, HeLa cells were selected due to its abundant expression of α-Syn, so the reduced α-Syn level could have been easily detected. Collectively, we present the first report of quercetin as a miR-7 enhancer and α-Syn inhibitor at low concentrations.

We hypothesized that quercetin inhibits α-Syn through the pri-miR-7/HuR pathway. However, we found similar inhibitive effects of quercetin in miR-7 KO HeLa cells, which means the upregulated miR-7 is not the major contributor of α-Syn suppression (Figure 6D). Indeed, quercetin has reduced α-Syn expression by around 60% in wildtype HeLa cells, and it would require more than 1,000-fold miR-7 upregulation to achieve a similar extent of α-Syn inhibition in the same cell line (Figure 1, Supplementary Figure 1A). Crucially, we found that α-Syn remained unchanged upon quercetin treatment when HuR was absent (Figure 6E, F). Nevertheless, the upregulation of miR-7 was still observed (Figure 6G). These results strongly imply that HuR is an essential part in the quercetin-mediated inhibition of α-Syn levels, although it seems not connected to the regulation of miR-7 biogenesis in our cellular system. Moreover, the biogenesis of miR-7 is also regulated by some other protein complexes, including SF2/ASF, NF45-NF90 and QKI (66–68). Therefore, the removal of HuR may allow alternative miR-7 regulation pathways to take over. This may explain why luteolin did not affect miR-7 level in wildtype HeLa, but significantly induced miR-7 in HuR KO HeLa cells.

HuR is known to stabilise target mRNAs through the AU-rich elements (AREs) in their 3’-UTRs (69). A recent study indicates that HuR interacts with the 3’-UTR of α-Syn mRNA and increases its stability in an mRNA decay assay in HeLa cells, although the knockdown of HuR only shows mild inhibition on α-Syn expression. Importantly, the stabilisation seems to be independent of miR-7 (70). In our study, quercetin significantly decreased α-Syn mRNA, and it is largely HuR-dependent. At the same time, in HeLa cells, the pri-miR-7/HuR pathway is not a major contributor of quercetin-mediated α-Syn inhibition (Figure 6H). Thus, quercetin may also directly interrupt HuR binding with the 3’-UTR of α-Syn mRNA, and destabilise the transcripts, contributing to the strong repression on α-Syn expression.

To summarise, we developed an on-bead screening platform, RP-CONA, for the identification of RNA-protein interaction modulators. We employed RP-CONA to look for inhibitors of pri-miR-7/HuR interactions. The most potent hit quercetin was proven to be a miR-7 enhancer as well as an α-Syn inhibitor, implying a potential as PD therapeutic. The mechanism of quercetin-mediated inhibition of α-Syn is HuR-dependent. Additionally, quercetin displayed positive effects on miR-7 levels, but the achieved upregulation was too small to directly affect α-Syn. Further work needs to be focused on the evaluation of quercetin in PD models, especially in the context of α-Syn overproduction and accumulation. In the meantime, the RP-CONA screening method can be used to identify more potent miR-7 enhancers. In summary, our work delivers a new platform for the identification of RNA-protein interaction modulators, and highlights quercetin as a potential hit compound towards treatment of PD.

## SUPPLEMENTAL MATERIAL

Supplemental material is available for this article.

## AUTHOR CONTRIBUTIONS

G.M. conceived the project. S.Z., M.A. and G.M. designed the work. S.Z. and S.R. performed the experiments and analysed the results. N.T.P. and J.K. helped optimising RP-CONA method. D.K. assisted in ImageXpress image analysis. G.M. coordinated the work and secured funding. S.Z., and G.M. wrote the manuscript with input from the other authors. All authors reviewed and approved the final manuscript.

## ACKNOWLEDGEMENTS

We would like to express our acknowledgement to Prof. Neil Carragher and Ashraff Makda for kindly providing us with the in-house compound library. We also thank James Longden for helping us with the ImageXpress service.

## ABBREVIATIONS

3’-UTR: 3 prime untranslated region
ATP: adenosine triphosphate
CPC: cetylpyridinium chloride
CTL: conserved terminal loop
CV: coefficient of variation
DHTS: dihydrotanshinone I
DNA: deoxyribonucleic acid
DTT: dithiothreitol
EDTA: ethylenediaminetetraacetic acid
EMSA: electrophoretic mobility gel shift assay
FITC: fluorescein isothiocyanate
FP: fluorescence polarisation
FRET: fluorescence resonance energy transfer
HCS: High Content Screening
HuR: human antigen R
KO: knockout
miR, miRNA: microRNA
mRNA: messenger RNA
MSI2: Musashi RNA binding protein 2
OA: oleic acid
PBS: phosphate-buffered saline
PCR: polymerase chain reaction
PD: Parkinson’s disease
PMSF: phenylmethanesulfonylfluoride
pre-miR, pre-miRNA: precursor microRNA pri-miR, pri-miRNA primary microRNA
qRT-PCR: real-time quantitative PCR
RBP: RNA binding protein
RNA: ribonucleic acid
Ro: Ro 08-2750
RP-CONA: RNA pull-down-Confocal Nanoscanning
RRM: RNA recognition motif
SD: standard deviation
SEM: standard error mean
shRNA: short hairpin RNA
siRNA: small interfering RNA
SNpc: substantia nigra pars compacta
TNF-α: tumour necrosis factor alpha
α-Syn: α-Synuclein

