## Supplementary Figures for "RNA pull-down-Confocal Nanoscanning (RP-CONA) detects quercetin as pri-miR-7/HuR interaction inhibitor that decreases α-Synuclein levels"

**Supplementary Figure 1**

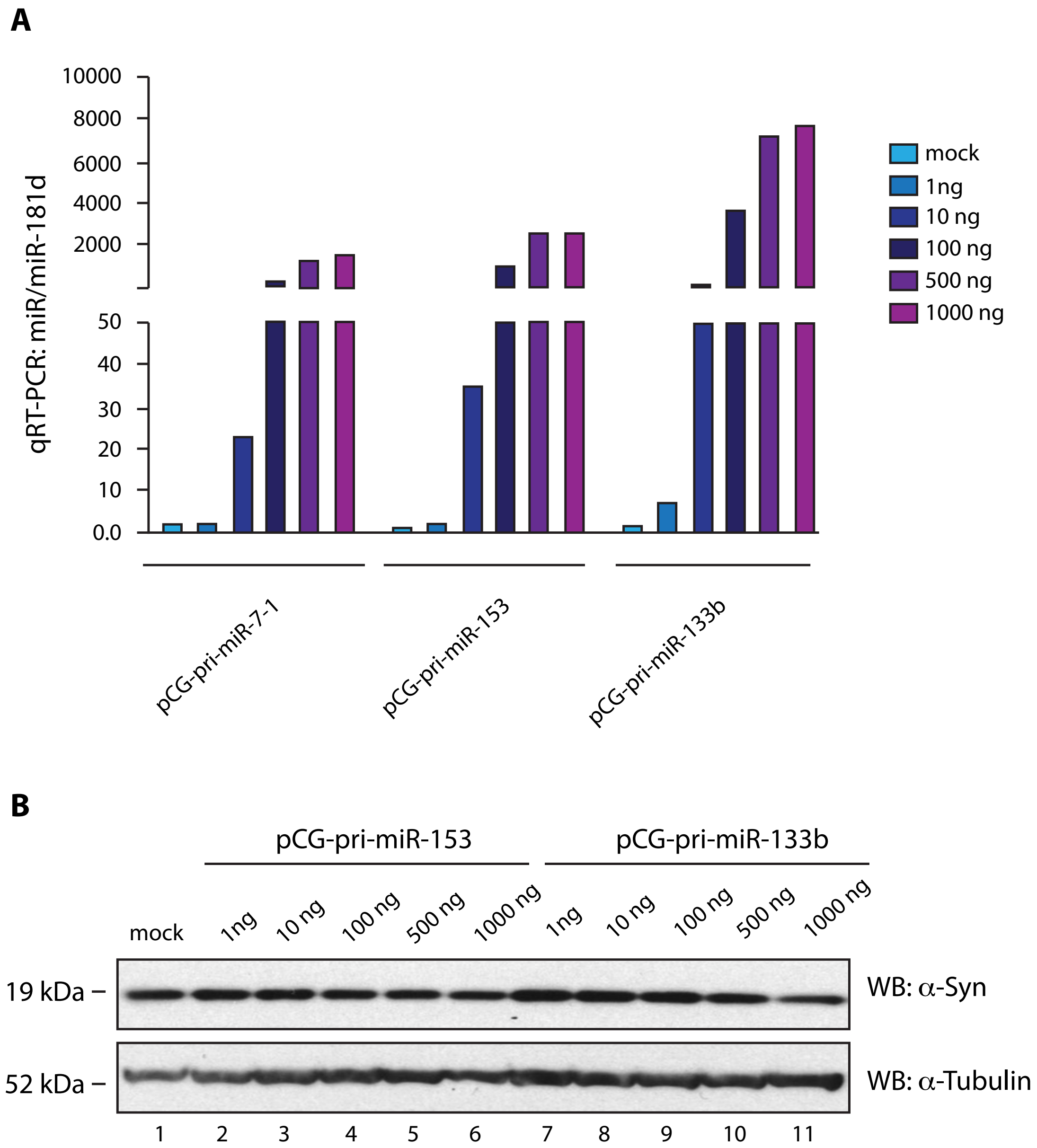


**Supplementary Figure 1.** MiR-7 is a major suppressor of α-Syn expression. (**A**) The upregulation of miRNAs after overexpression. The mature miR-7, miR-153 and miR-133b levels from HeLa cells transfected with increasing amount of corresponding pCG plasmids were determined by qRT-PCR and normalised to mock (no DNA transfected). (**B**) MiR-153 and miR-133b exert mild inhibition on α-Syn expression.1: Mock HeLa cells without DNA transfected. 2-6: Increased amount of pCG-pri-miR-153 wase transfected into HeLa cells. 7-11: Increased amount of pCG-pri-miR-133b was transfected into HeLa cells. The expression of α-Syn and α-tubulin were detected by western blot 48 hours after transfection.

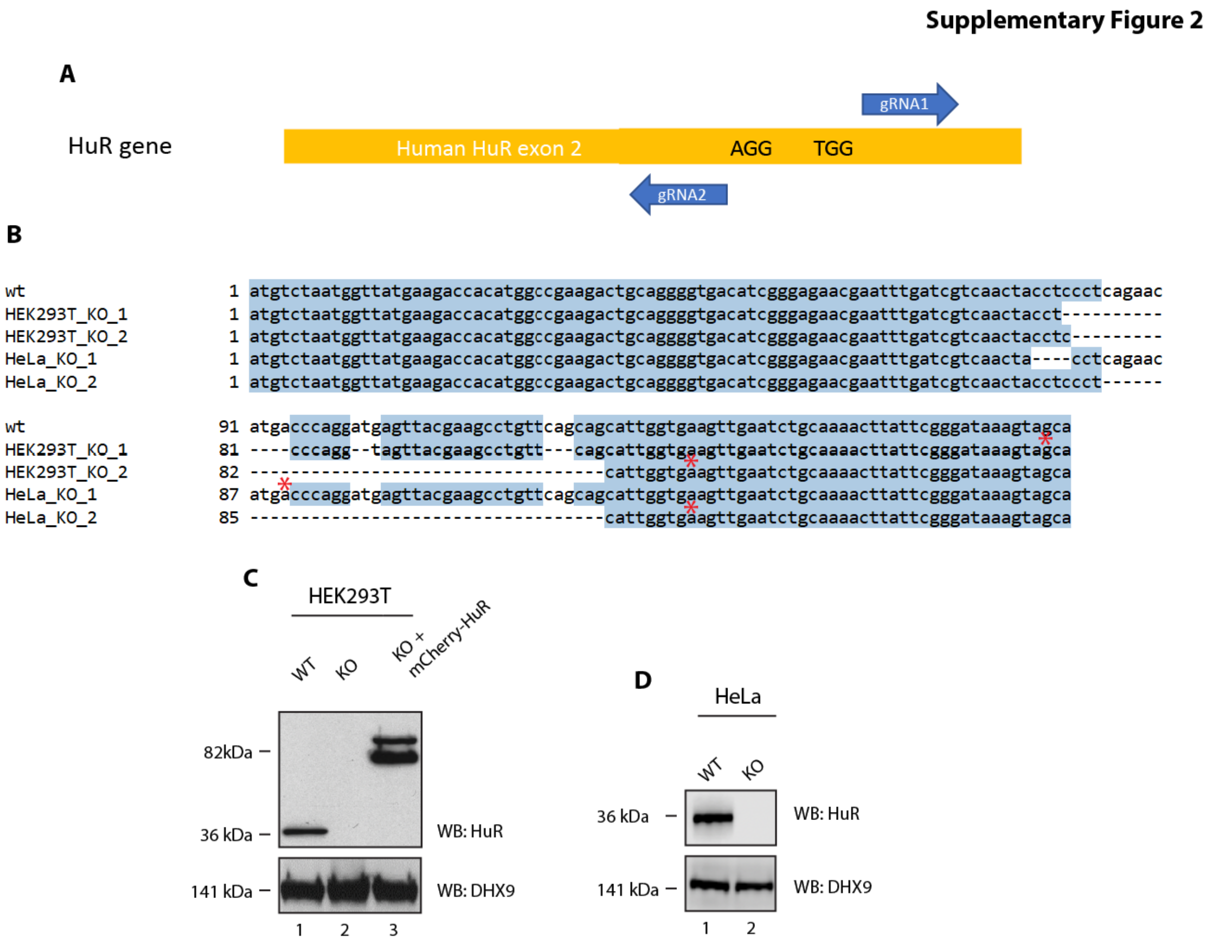


**Supplementary Figure 2.** HuR knockout and overexpression. (**A**) Design of HuR knockout by CRISPR-Cas9. Human HuR exon2 is shown in orange. A pair of guide RNAs are shown in blue arrows. PAM sequences are presented. (**B**) Sequence alignment of HuR knockouts. The DNA sequences of HuR exon2 of HEK293T and HeLa knockouts were aligned to the wildtype sequence in Clone Manager 10. Termination signals resulting from frame shifts are annotated by red stars. (**C**) Validation of HuR knockout and mCherry-HuR overexpression in HEK293T cells. The endogenous HuR and overexpressed mCherry-HuR levels were tested against HuR antibody in western blot. DHX9 was tested as a reference protein. 1: Wildtype HEK293T. 2: HuR KO HEK293T. 3: HuR KO HEK293T transfected with pJW99-HuR. (**D**) Validation of HuR knockout in HeLa cells. The HuR levels were tested against HuR antibody in western blot. DHX9 was tested as a reference protein. 1: Wildtype HeLa. 2: HuR KO HeLa.

### Supplementary Figure 3

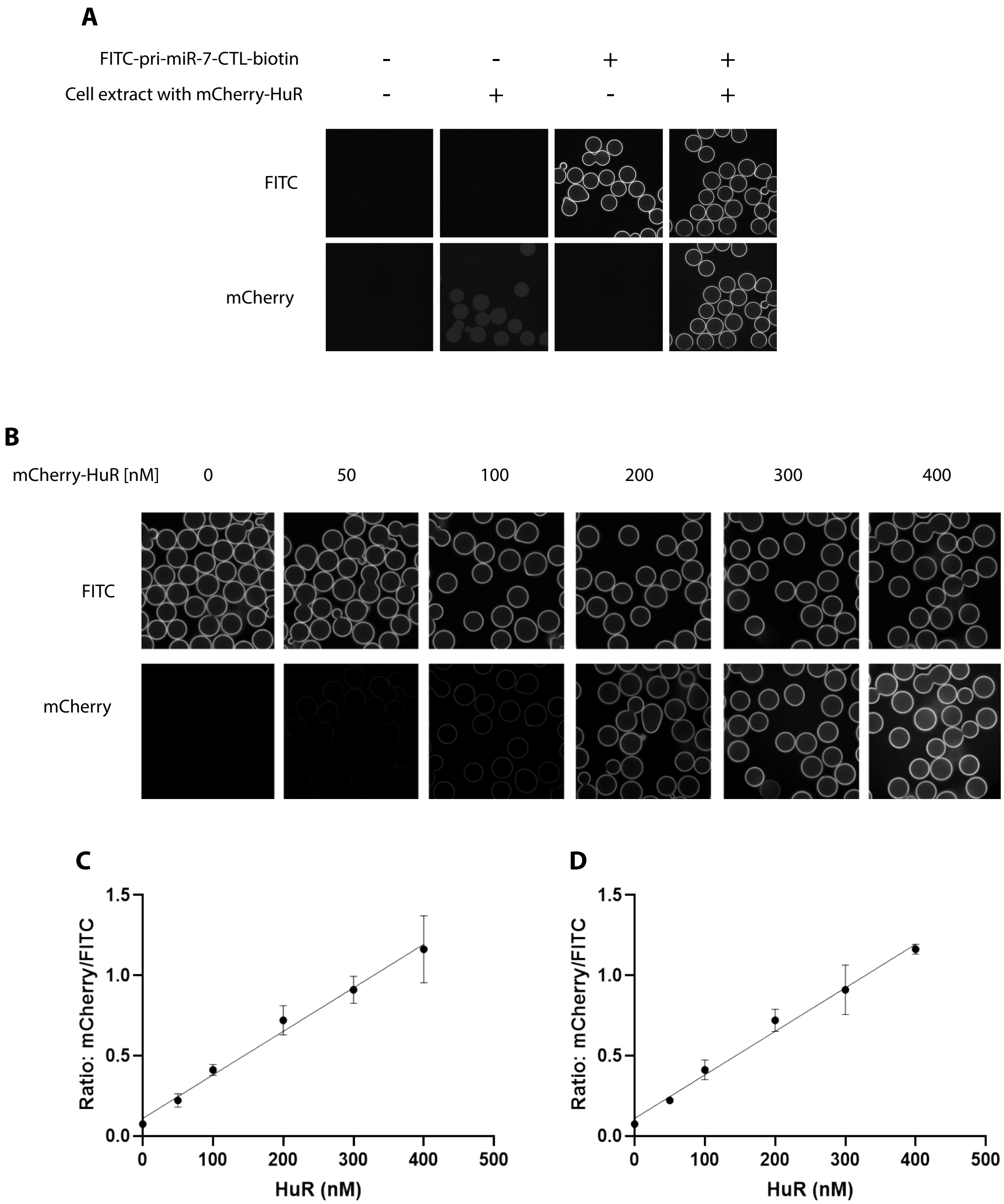


**Supplementary Figure 3**. Basic responses of RP-CONA in ImageXpress. (**A**) Beads images of blank beads; blank beads incubated with cell lysates containing mCherry-HuR; FITC-pri-miR-7-CTL-beads incubated in lysates-free buffer; and FITC-pri-miR-7-CTL-beads after mCherry-HuR pulldown. (**B, C, D**) RP-CONA senses the increase of mCherry-HuR. FITC-pri-miR-7-CTL-beads were incubated with cell lysates containing an increased concentration of mCherry-HuR. All the quantifications were obtained from three technical repeats. Mean mCherry/FITC ring intensities and maximum SD between the beads in each well (**C**), or SD between triplicates (**D**) are shown.

**Supplementary Figure 4**

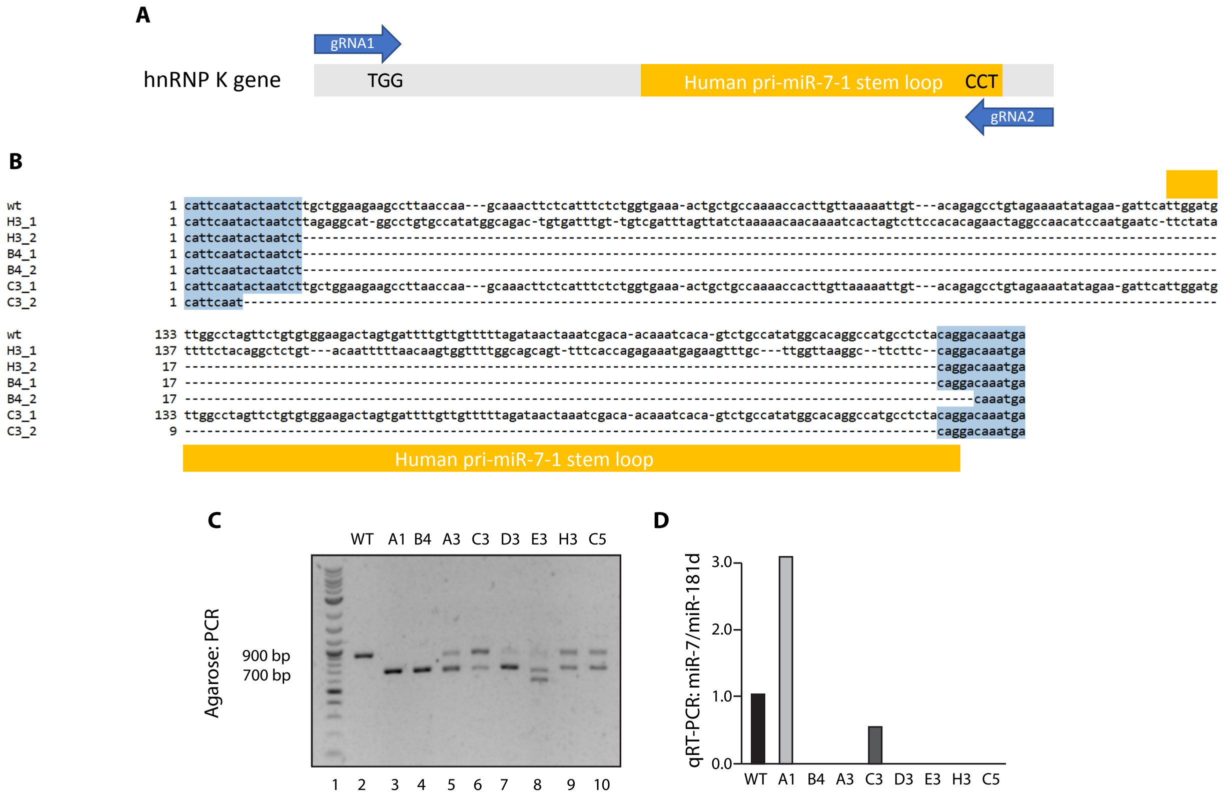


**Supplementary Figure 4.** MiR-7 knockout by CRISPR-Cas9. (**A**) Design of miR-7 knockout by CRISPR-Cas9. Human pri-miR-7-1 stem loop sequence is shown in orange. Other Regions are in grey. A pair of guide RNAs flanking the pri-miR-7-1 stem loop gene are shown in blue arrows. PAM sequences are presented. (**B**) Sequence alignment of miR-7 knockouts. Sequencing results of miR-7 knockouts were aligned to the same region of wildtype HeLa gene in Clone Manager 10. Regions originally encoding pri-miR-7-1 stem loop was indicated. B4 is a miR-7$^{-/-}$ as the miR-7 stem loop was deleted in both alleles. C3 is a miR-7$^{+/-}$ as miR-7 was deleted in one allele while the other allele is intact. H3 has miR-7 deleted in one allele. The other allele, however, the miR-7 stem loop was re-assembled in an opposite orientation. Therefore, H3 is a miR-7$^{-/-}$. (**C**) Genotyping and characterisation of miR-7 KO HeLa. Genomic DNA was PCR amplified using a pair of primers flanking pri-miR-7-1 stem loop. The deleted stem loop region was expected to be around 200 bp. Bands around 700 bp represent potential KO. (**D**) MiR-7 level of miR-7 KO HeLa. Mature miR-7 and miR-181d levels of wildtype HeLa or miR-7 knockouts were determined by qRT-PCR.
